# Database-integrated genome screening (DIGS): exploring genomes heuristically using sequence similarity search tools and a relational database

**DOI:** 10.1101/246835

**Authors:** Henan Zhu, Tristan Dennis, Joseph Hughes, Robert J. Gifford

**Author notes:** These authors contributed equally to this work.

## Abstract

A significant fraction of most genomes is comprised of DNA sequences that have been incompletely investigated. This genomic ‘dark matter’ contains a wealth of useful biological information that can be recovered by systematically screening genomes *in silico* using sequence similarity search tools. Specialized computational tools are required to implement these screens efficiently. Here, we describe the database-integrated genome-screening (DIGS) tool: a computational framework for performing these investigations. To demonstrate, we screen mammalian genomes for endogenous viral elements (EVEs) derived from the Filoviridae, Parvoviridae, Circoviridae and Bornaviridae families, identifying numerous novel elements in addition to those that have been described previously. The DIGS tool provides a simple, robust framework for implementing a broad range of heuristic, sequence analysis-based explorations of genomic diversity.

**Availability:** http://giffordlabcvr.github.io/DIGS-tool/

**Supplementary information:** Supplementary data are available at Bioinformatics online.

## INTRODUCTION

A large fraction of most published genome assemblies consists of DNA sequences that have no known function. The diverse assortment of pseudogenes, transposons, and non-coding DNA elements that make up this genomic ‘dark matter’ contain useful biological information, and can reveal important insights into the evolution of genes and genomes. Sequence similarity search tools, such as the *Basic Local Alignment Search Tool* (BLAST) (Altschul, et al., 1997), provide powerful devices for investigating this poorly characterized component of genomes *in silico*.

The BLAST program takes a protein or nucleic acid sequence (the ‘probe’ or ‘query’) as input, and efficiently searches target databases for sequences exhibiting local similarity to this query (‘hits’). Results are reported in the form of a ranked list of hits, each associated with a statistical significance. Genomic diversity can be explored by using BLAST to systematically survey genome databases for specific sequences of interest. We have developed a variation of this general approach, called ‘database-integrated genome screening’ (DIGS), in which a relational database is used to record which searches have been performed, and to capture their results. This facilitates the implementation of automated screens that can be performed on a large scale, and provides a powerful, well-established framework for interrogating and manipulating the data they produce. In addition, it provides all the benefits of a relational database management system (RDBMS) with respect to features such as data recoverability, multi-user support and network access.

Here we describe the DIGS tool, an open source PERL program for implementing DIGS. We use an example based on endogenous viral elements (EVEs) to demonstrate the functionality and features of the DIGS tool.

## FUNCTIONALITY

The DIGS tool is written in PERL. It uses the BLAST program to perform sequence similarity searches, and MySQL as an RDBMS. All of these programs are cross-platform and freely available for non-commercial use. The DIGS tool itself is available as a public repository. The repository includes a detailed guide to installing and running the program (http://giffordlabcvr.github.io/DIGS-tool/).

As illustrated in Figure 1, the screening process entails performing two distinct BLAST-based steps for each probe-target pairing. In the first step, the probe sequence is used to search the target database. In the second, hits from the first BLAST search are extracted from target databases and then genotyped via BLAST comparison to the reference sequence library. Thus, in this second step, the extracted sequences are used as probes, and the reference sequence library as the target database. For each extracted sequence, the best hit from this second BLAST search is taken as the ‘genotype’. A genotyping step is included because probes used in similarity search based screens will often yield hits to a wide range of related sequences, and it is important to be able to distinguish between these different kinds of hits. For example, consider a gene that has two paralogs, ‘X’ and ‘Y’, where screening with a probe of type X yields hits to both X and Y. Genotyping these hits by comparing each to a library of representative reference sequences that includes both X and Y allows the different types of hit to be differentiated. While such BLAST-based genotyping clearly cannot substitute for in-depth phylogenetic analysis, it is an efficient approach for broad classification of sequence diversity (Gifford, et al., 2006).

**Figure 1.**
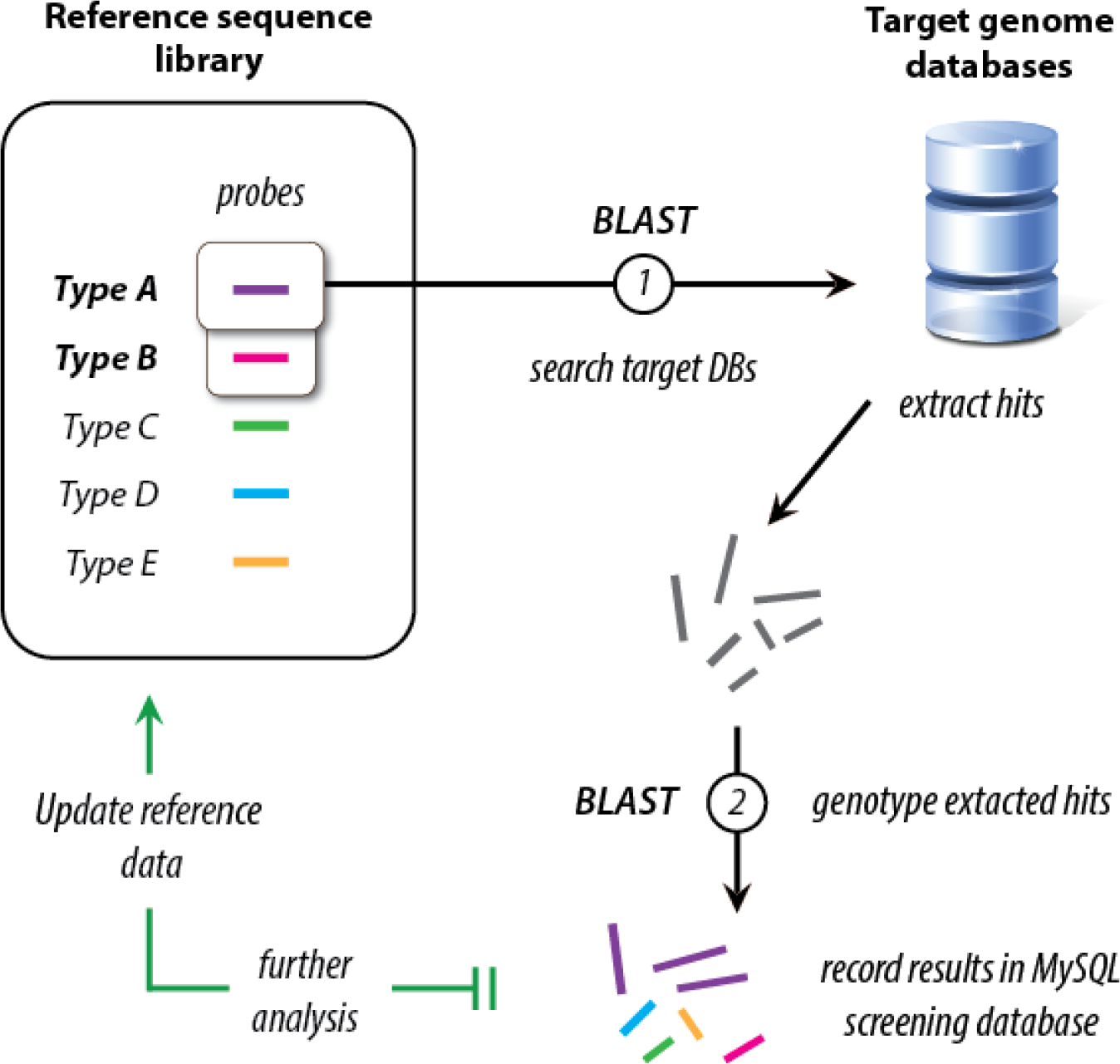
Genome screening using the database integrated genome screening (DIGS) tool. The DIGS tool is a project-oriented framework. Each DIGS project consists of: (i) a set of ‘target’ sequence databases (e.g. genome assemblies of a chosen target species or species group); (ii) a library of reference sequences for the gene(s) or DNA element(s) under investigation; (iii) a set of query sequences (‘probes’) for use in screening (usually a subset of sequences from the reference library); (iv) a relational database comprising three core tables. Black arrows show the main DIGS program loop. A probe is used to search a specified target database(s) (BLAST 1). All sequences producing statistically significant matches to this probe are extracted and genotyped by comparison to the reference sequence library (BLAST 2). A MySQL relational database records which screens have been performed, and their results. Iterative DIGS workflows can be developed, wherein sequences recovered by screening are analyzed (e.g. phylogenetically), and information derived from this analysis is used to incrementally update the reference sequence library (green arrows). BLAST 2 can then be repeated for all extracted sequences, using the updated reference library. In addition, new BLAST 1 searchers can be performed using newly defined sequences as probes. By iteratively expanding the reference library in this way the accuracy and sensitivity of screens can be progressively refined (see Table 1).

To illustrate how the DIGS tool can be used, we mined vertebrate genomes for nonretroviral endogenous viral elements (EVEs). An EVE is a DNA sequence derived from a virus that is present within the germ line of another organism. All EVEs were discovered based on their similarity in sequence to extant viruses (Patel, et al., 2011), and in the post-genomic era, these discoveries have increasingly been made *in silico* (Johnson, 2015).

In vertebrate genomes, the vast majority of EVEs are derived from retroviruses (family *Retroviridae*). However, progress in whole genome sequencing has revealed that sequences derived from other virus families are also present, albeit at a much lower frequency. We used the DIGS tool to search for these rare, non-retroviral EVEs in vertebrate genomes. We obtained a reference sequence library from the NCBI virus genomes database, comprising 53,610 polypeptide gene products derived from a total of 4,297 viruses. Representative sequences from this library were selected to use as probes for screening (**Table S1**). Probes were chosen to represent genetic diversity within five virus families (*Bornaviridae*, *Filoviridae*, *Circoviridae*, *Parvoviridae* and *Hepadnaviridae*) that have been shown to occur as EVEs in vertebrate genomes (Katzourakis and Gifford, 2010). Polypeptide probes were used to screen the genomes of 187 vertebrate species (**Table S2**) *in silico* for EVEs derived from these five families.

Initial screening generated numerous hits that were revealed on inspection not to be *bona fide* non-retroviral EVEs, but spurious matches to endogenous retroviruses (ERVs) and transposons. We therefore incorporated representatives of these sequences into the reference sequence library, and repeated the genotyping step for all the loci matched by our screen. By iteratively updating in this way (see Figure 1), we eliminated false positive hits from our final screening output (Table 1).

**Table 1.** Summary of 1111 vertebrate EVEs identified using the DIGS tool.

| Virus family | Vertebrate lineage |  |  |  |  |  |
| --- | --- | --- | --- | --- | --- | --- |
|  | *Fishes* |  | *Squamates* |  | *Mammals* |  |
|  | **S** | **NS** | **S** | **NS** | **S** | **NS** |
| **ssRNA** |  |  |  |  |  |  |
| *Arenaviridae* | 0 (7) |  | 0 (27) |  |  |  |
| *Flaviviridae* |  |  |  |  |  | 0 (1) |
| *Bornaviridae* | **1** | **2** | **8** | **1** | **265** | **98 (98)** |
| *Filoviridae* | **2 (30)** |  | **2 (88)** |  | **37 (51)** | **17 (17)** |
| *Paramyxoviridae* |  | **2** |  |  |  |  |
| *Nyamiviridae* |  | **1** |  |  |  |  |
| **ssDNA** |  |  |  |  |  |  |
| *Circoviridae* |  | **3** |  | **4** |  | **58** |
| *Parvoviridae* | **3** | **7** | **14** | **11** | **152** | **182** |
| **Retro-transcribing** |  |  |  |  |  |  |
| *Hepadnaviridae* |  | 0 (2) | **48** | **193 (195)** |  | 0 (1) |
| *Caulimoviridae* |  | 0 (88) |  | 0 (628) |  | 0 (1) |
| **Totals** | **6** | **15** | **72** | **209** | **454** | **355** |

**Legend:** S=number of EVEs matching derived from structural proteins; NS=number of EVEs derived from non-structural proteins. Numbers in brackets to the right show the number of hits obtained in the initial DIGS screen (if different from the final count). Bold numbers to the left show the final count, following updates to the reference library to include endogenous retrovirus (ERV), retrotransposons, and genomic sequences that produced spurious hits to non-retroviral EVEs.

The relational database component of the DIGS tool provides a powerful framework for interrogating data produced by screening. Furthermore, extending the database to include additional ‘side-data’ for (i) the target species/databases being screened, and/or (ii) sequences in the reference library, allows screening output to be summarized based on these data as well. For example, here we created a database table containing additional taxonomic data for the vertebrate species we screened, and another for the virus genes that were represented as EVEs in vertebrate genomes, indicating which encoded structural versus non-structural proteins. This allowed us to summarize the results of screening according to these criteria (see Table 1). The database generated by this screen, the SQL commands used to generate Table 1, and the associated DIGS project, are available at:

## CONCLUSIONS

Molecular sequence data are highly information rich, and are accumulating much faster than they can be analysed. Accordingly, there is currently unprecedented scope for new biological discoveries to be made by heuristically exploring genomic diversity *in silico*. The DIGS tool can provide a robust and adapable framework for implementing a broad range of such investigations.

## Acknowledgements

We thank Andrew Davison, Jamie Henzy, Pablo Murcia and Sam Wilson for helpful discussions and input on the manuscript.

